# Enhanced anti-tumor effects through continuous administration of engineered CAR-macrophages derived from pluripotent stem cell-derived myeloid cell lines

**DOI:** 10.1101/2024.07.22.604686

**Authors:** Yuya Atsumi, Akira Niwa, Tatsuro Kumaki, Shigeki Yagyu, Yozo Nakazawa, Megumu K. Saito

**Affiliations:** Department of Clinical Application, Center for iPS cell Research and Application, Kyoto University, Kyoto, Japan; Department of Pediatrics, Graduate School of Medical Science, Kyoto Prefectural University of Medicine, Kyoto, Japan; Center for Advanced Research of Gene and Cell Therapy in Shinshu University (CARS), Shinshu University School of Medicine, Matsumoto, Japan; Department of Pediatrics, Shinshu University School of Medicine, Matsumoto, Japan

**Keywords:** CAR, Macrophage, Induced pluripotent stem cell, Therapy, Cost

## Abstract

Even after chimeric antigen receptor (CAR)-based immunotherapy has dramatically changed therapeutic approaches for malignancies, balancing therapeutic efficacy with labor and financial cost remains a major problem for immunotherapy. Current study developed a cost-effective and enhanced approach to chimeric antigen receptor (CAR)-macrophage therapy for cancer and demonstrated its therapeutic effects by repeated administration of anti-HER2 CAR macrophages generated from human pluripotent stem cell (PSC)-derived immortalized myeloid cell lines (ML). These ML-derived CAR macrophages (CAR-ML-MPs) exhibit potent antigen-specific killing activity against HER2-expressing tumor cells by phagocytosis *in vitro* and effectively inhibit tumor progression *in vivo*, which is enhanced by repeated administration. CAR-ML-MPs provide a promising off-the-shelf cellular resource for tumor adoptive cell immunotherapy, solving the cost and time problems associated with conventional CAR-based immunotherapy.

## Introduction

Even after chimeric antigen receptor (CAR)-based immunotherapy has dramatically changed therapeutic approaches for malignancies, balancing therapeutic efficacy with labor and financial cost remains a major problem for immunotherapy^1^. Immune cells used in adoptive cellular immunotherapy (ACT), such as T cells, are usually collected from the peripheral blood of patients with cancer using apheresis^2^. However, the burden of apheresis is high for patients; approximately 5–10% of failure cases due to poor lymphocyte condition and counts^2^. Moreover, ACT is extremely expensive, owing to the need for personalized medicine. Tisagenlecleucel (Kymriah, Novartis), the first-in-class CAR-T therapy approved by the United States Food and Drug Administration in August 2017, costs more than $350,000 per treatment^1^. Given that CAR-T therapy against CD19-positive B-cell malignancies relapsed in approximately 36% of cases^3^, the cost of consecutive treatments would be even higher. Consequently, off-the-shelf products that can be readily available to patients and healthcare systems are required.

Pluripotent stem cells (PSCs), including embryonic stem cells (ESCs) and induced pluripotent stem cells (iPSCs), are ideal sources for immune cell production owing to their proliferative capacity, pluripotency, and accessibility to genetic manipulation. Therefore, CAR-expressing PSC-derived immune cells, such as T cells, natural killer cells, and macrophages, have been developed for broader use in CAR technology. CAR expression can enhance the anti-tumor activity of PSC-derived immune cells^4^. In particular, CAR macrophages have attracted increasing attention in immunotherapy for intrinsic properties of macrophages, such as their immunomodulatory capacity and ability to infiltrate and phagocytose tumor cells^5^. Additionally, CAR macrophages are expected to function as specialized antigen-presenting cells (APCs) and initiate long-term adaptive immunity against tumors^6^, in addition to act as direct tumor attack cells^6^. However, differentiation of PSCs into immune cells is expensive and laborious. Moreover, the differentiation process takes more than a month, and the number of cells obtained is limited^4,7,8^.

PSC-derived myeloid cell line (ML) is an immortalized monocytic cell line capable of continuous expansion through M-CSF and GM-CSF, with the ability to differentiate into macrophages (ML-MPs) and dendritic cells^9^. Use of ML-MPs enables immediate preparation of macrophages for ACT and could be an alternative cell source. In this study, we aimed to demonstrate *in vitro* cytotoxic potential of ML-derived Doxycycline (DOX)-inducible anti-HER2 CAR macrophages (CAR-ML-MPs) and their ability to reduce tumor burden and prolong overall survival in a tumor-bearing mouse model. In addition, we further aimed to show that effects were enhanced when ML-MPs were administered continuously, taking advantage of less laborious propagation and preparation.

## Materials and methods

### Study approval

The use of human ESCs was approved by the Ministry of Education Culture, Sports, Science and Technology of Japan (MEXT). The study plan for recombinant DNA research was approved by the recombinant DNA experiments safety committee of Kyoto University. Animal studies were approved by the institutional review board in the Center for iPS Cell Research and Application, Kyoto University (#21-149-3). All methods were performed in accordance with the relevant guidelines and regulations.

### Cell lines and vectors

A human iPSC line 409B2 and a human ESC line KhES1 were kindly provided by Dr. Shinya Yamanaka (Kyoto University, Japan) and Dr. Norio Nakatsuji (Kyoto University, Kyoto, Japan), respectively. A luciferase-expressing leukemic cell line Luc-K562^10^ was kindly provided by Dr. Yasushi Uemura (National Cancer Center, Tokyo, Japan). A plasmid vector encoding anti-HER2 CAR construct was kindly provided by Dr. Cliona M. Rooney (Baylor College of Medicine)^11^. We generated DOX-inducible anti-HER2 CAR construct by inserting anti-HER2 CAR under the tetracycline-response element (TRE). CAR-PSCs were established by introducing this construct by electroporation (NEPA21, Nepa Gene, Chiba, Japan). A human breast cancer cell line, OCUB-F^12^ was purchased from RIKEN BioResource Research Center (Tsukuba, Ibaraki, Japan). Luciferase and HER2-expressing K562 cells (Luc-HER2-K562) were established by introducing the luciferase gene and the extracellular domain of the HER2 gene by electroporation. Luc-GFP-OCUB-F was established by introducing the luciferase and GFP genes by lentivirus vector.

### Transfection

The introduction of CAR into iPS cells and the introduction of the HER2 gene into tumor cells were both performed by mixing 1×10^5^ cells with 5 ng of PiggyBac plasmid and 5 ng of transposase expression plasmid. The transduction was carried out under conditions optimized and prepared in advance by the manufacturer of the electroporation device.

### Cell culture

hPSCs and hESCs were cultured on tissue culture dishes coated with Laminin511-E8 (#892021; Nippi, Osaka, Japan) in StemFit medium (Ajinomoto, Tokyo, Japan) as previously described^13,14^. K562 was cultured in RPMI-1640 medium (#30264-56; Nacalai Tesque, Kyoto, Japan) containing 10% fetal bovine serum (FBS, #10438026, Gibco, New York, USA). OCUB-F was cultured in DMEM medium (#08459-64; Nacalai Tesque) containing 10% FBS.

### Establishment of MLs

Hematopoietic differentiation from human PSC lines into monocytic cells was performed as previously reported^8^. MLs were established by lentiviral transduction of three genes, cMYC, MDM2, and BMI1, into monocytic cells^9^. MLs were cultured in α-MEM (Nacalai Tesque) supplemented with 20 % FBS, 50 ng/mL recombinant human macrophage colony-stimulating factor (#216-MC; M-CSF, R&D Systems, Minneapolis, MN, USA) and 50 ng/mL recombinant human granulocyte-macrophage colony-stimulating factor (#215-GM; GM-CSF, R&D Systems). For macrophage differentiation, MLs were cultured in RPMI-1640 medium containing 10 % FBS and 100 ng/µl M-CSF for 7 days and adherent cells were collected as macrophages. For the induction of CAR, DOX (final concentration: 1 µg/ml) was added at the start of macrophage induction and was continuously administered during subsequent media changes. Alternatively, dimethyl sulfoxide (DMSO, #D2770; Sigma-Aldrich St. Louis, MO, USA, final concentration; 0.1 %) was added as control.

### May-Giemsa staining

Cytology specimens of ML-MPs were prepared using Shandon Cytospin 4 Cytocentrifuge (Thermo Fisher Scientific, Sunnyvale, CA, USA), and then May-Giemsa staining (#HX85293624, #HX082310; Merck, Darmstadt, Germany) was performed according to manufacturer’s protocol.

### Flow cytometry

Flow cytometric analysis data were collected using BD FACSAria II (BD bioscience, San Jose, CA, USA) and subsequently analyzed with the FlowJo software package (Flowjo LLC, Ashland, OR, USA). Dead cells were removed by 4,6-diamidino-2-phenylindole (DAPI) staining (#D9542; Sigma-Aldrich). The following antibodies were used in this study flow cytometry analysis: CD36-FITC (#IM0766U; Beckman Coulter, Brea, CA, USA), CD45-FITC (#368508; BioLegend, San Diego, CA, USA), HLA-DR-FITC (#307604; Bio Legend), CD206-FITC (#551135; BD bioscience), CD86-FITC (#555657; BD bioscience), CD11b-Alexa488 (#557701; BD bioscience), CD340(erbB2/HER-2)-PE (#324406; Bio Legend), CD14-APC (#IM2580; Beckman Coulter), CD11c-APC (559877; BD bioscience), Fc Receptor Blocking Solution (#422302; Bio Legend), FITC IgG1 isotype control (#400110; Bio Legend), FITC IgG2a isotype control (#349051; BD bioscience), PE IgG1 isotype control (#400112; Bio Legend), APC IgG1 isotype control (#400122; Bio Legend), and APC IgG2a isotype control (#400512; Bio Legend). For confirmation of anti-HER2 CAR expression, primary staining was performed with human HER2/ERBB2 Protein-His tag (#10004H08H100; Sino Biological, Beijing, China), followed by secondary staining with anti-His Tag APC (#IC050A; R&D Systems).

### Cytotoxicity assay *in vitro*

Luc-HER2-K562 and Luc-GFP-OCUB-F and ML-MPs were co-cultured in 96-well plates. Based on preliminary studies, the effector/target ratio was set to 5:1 for ML-MPs and Luc-HER2-K562, and 3:1 for ML-MPs and Luc-GFP-OCUB-F. As for the actual number of cells, 5×10^3^ cells of LUC-HER2-K562 was co-cultured with 2.5×10^4^ cells of ML-MPs, and 7.5×10^3^ cells of Luc-GFP-OCUB-F was co-cultured with 2.25×10^4^ cells of ML-MPs in 200 µl of RPMI-1640 with 10% FBS. After 3 days of incubation, 50 µl of PicaGene MelioraStar-LT luminescence reagent (#MLT100; Toyo B-Net, Tokyo, Japan) was added and cultured additional 5 minutes. Bioluminescence was measured with EnVision 2104-multilabel reader (PerkinElmer, Waltham, MA, USA).

### Time-lapse imaging

ML-MPs were stained with PKH26 Red Fluorescent Cell Linker Kit for Phagocytic Cell Labeling (#PKH26PCL-1KT, Sigma-Aldrich). 2 x 10^4^ cells of ML-MPs were co-cultured with 2 x 10^4^ cells of Luc-GFP-OCUB-F in 110 µl of RPMI-1640 with 10% FBS and 100 ng/ml M-CSF and 1 µg/ml DOX. Time-lapse imaging data were acquired using FV1000 confocal microscopy (Olympus, Tokyo, Japan).

### Cytotoxicity assay *in vivo*

Luc-HER2-K562 (1×10^5^ cells) and ML-MPs (1×10^6^ cells) were suspended in 200 µl of RPMI-1640 with 100 ng/ml of M-CSF and 20% of Matrigel (#354248; Corning, Corning, NY, USA). The mixture was transplanted subcutaneously into NOG mice (Ito et al., 2002) (Central Institute for Experimental Animals). In order to induce CAR expression in mice, aqueous solutions including 1 mg/ml Doxycycline hyclate (#AA22608; Biosynth, Berkshire, UK), 50 mg/ml sucrose (#30403-55; Nacalai Tesque), and 72 µg/ml Sulfamethoxazole and trimethoprim (Shionogi, Osaka, Japan) were administered as drinking water. In the sequential administration regimens, ML-MPs were administered by subcutaneous injection of 2 x10^6^ cells into the tumor inoculation site every day for 6 days, starting the day after the initial treatment with the same number of cells as in the single administration. For the measurement of luciferase activity, 150 mg/kg of Luciferin (#XLF-1; SPI, Tokyo, Japan) was injected subcutaneously (Figure 2) or intraperitoneally (Figure 4) into mice and bioluminescence imaging was performed using IVIS Lumina series III (PerkinElmer). During the measurement, sedation was performed by intraperitoneal injection of a mixture of medetomidine (0.3 mg/ kg), midazolam (3 mg/kg), and butorphanol (5 mg/kg).

### Statistical analysis

Statistical analysis of all data was performed using GraphPad Prism version 6 (GraphPad Software, San Diego, CA, USA). Dunnett’s test was performed as *a priori* multiple comparisons *in vitro* and *in vivo* assays. Log-rank (Mantel-Cox) test was performed for the comparison of survival curves. p < 0.05 was defined as a significant difference.

### Data Availability

The data that support the findings of this study are available in the methods and/or supplementary material of this article and are available on request from the corresponding author.

## Results

### Establishment of doxycycline-inducible CAR-myeloid cell line derived from human pluripotent stem cells

We prepared two human PSC strains (iPSC 409B2 and KhES1) transduced with a Tet-inducible anti-HER2 CAR^11^ expression system (Figure 1A). After differentiation into monocytes, MLs (CAR-MLs) were established by transducing cells with BMI1, c-MYC, and MDM2^9^. CAR-MLs readily proliferated in the presence of M-CSF and GM-CSF for longer than two months (Figure 1B) and differentiated into macrophages (CAR-ML-MPs); Once they differentiated into macrophages, their morphology changed distinctively (Figure 1C), and cell division ceased. They expressed not only CAR but also appropriate cell surface markers, including CD45 (a pan-leukocyte marker) and CD14 and CD11b (markers for monocytes and macrophages). Additionally, macrophages are generally classified into pro-inflammatory M1 macrophages and anti-inflammatory M2 macrophages. In this regard, they were positive for both M1 markers (CD11c, HLA-DR, and CD86) and M2 markers (CD206 and CD36) (Figures 1D and 1E). Furthermore, RNA expression analysis showed that several functional factors, such as OX40L and 4-1BBL, which are costimulatory molecules involved in the activation and survival of immune cells, are expressed at low levels (Figure 1F). Given these findings, we believe that CAR-MLs can serve as a useful cell source for off-the-shelf CAR macrophage products.

**Figure 1.**
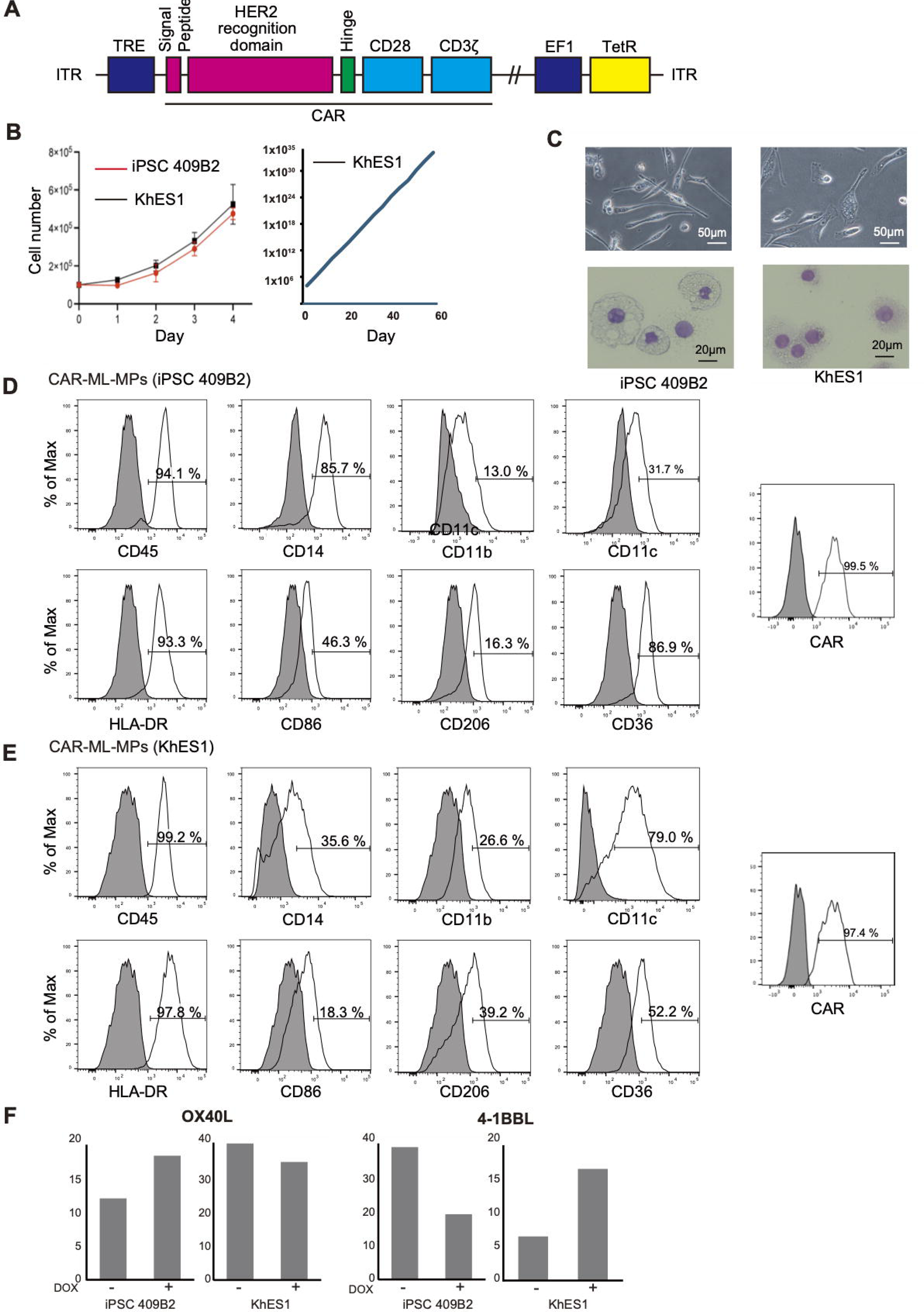
Generation of anti-HER2 CAR transduced PSC-MLs and ML-MPs. (A) Schematic diagram of DOX-inducible anti-HER2 CAR construct. The construct includes HER2 recognition domain, intracellular hinge, CD28 and CD3ζ. TRE: tetracycline-response element. EF1: EF1 Alpha Promoter. TetR: Tet Repressor. ITR: inverted terminal repeat. (B) Proliferation curve of PSC-MLs derived from CAR-PSCs. On the left is mean ± standard deviation (SD) data for one passaging culture (n = 3, independent experiment). On the right, growth curves of KhES1-derived MLs cultured for 60 days. (C) Morphology of ML-MPs. Phase-contrast images (Upper panels) and May–Giemsa staining images (Lower panels). (D, E) Representative histograms showing macrophage cell-surface markers on CAR-ML-MPs. Panels (D) and (E) depict data from iPSC 409B2 and KhES1 strains, respectively. The rightmost panel displays histograms showing anti-HER2 CAR expression on the surface of CAR-ML-MPs in the presence of DOX. Gray fills represent isotype controls in all panels. (F) Expression of OX40L (left two panels) and 4-1BBL (right two panels). The relative quantity (Rq) of RNA expressions was measured in ML-MPs derived from the iPSC 409B2 strain (left) and the KhES1 strain (right) under both conditions, with and without DOX, compared to undifferentiated cells.

### CAR-ML-MPs have anti-tumor ability *in vitro*

To evaluate their anti-tumor ability, we co-cultured CAR-ML-MPs with HER2- and luciferase-expressing tumor cell lines (Luc-HER2-K562 and OCUB-F) (Figure 2A). CAR-ML-MPs showed significant cytotoxicity against both Luc-HER2-K562 (p=0.0481 and p=0.0072 in KhES1 and 409 B2 strains, respectively) (Figure 2B) and OCUB-F (p=0.0200 and p=0.0288 in KhES1 and 409 B2 strains, respectively) (Figure 2C) cells. Under certain conditions, the antitumor effect of CAR-ML-MPs was observed even in the absence of HER2 expression on tumor cells (Figure 2D). This suggests that ML-MPs possess intrinsic cytotoxic activity independent of CAR. However, significant differences between the presence and absence of CAR expression were only observed when targeting HER2-expressing tumor cells. Notably, time-lapse imaging confirmed that CAR-ML-MPs phagocytosed tumor cells (Figure 2D and Supplemental Movie 1). CAR-ML-MPs have therefore cytotoxic ability against tumor cells, which is at least partially exerted by direct phagocytosis of tumor cells.

**Figure 2.**
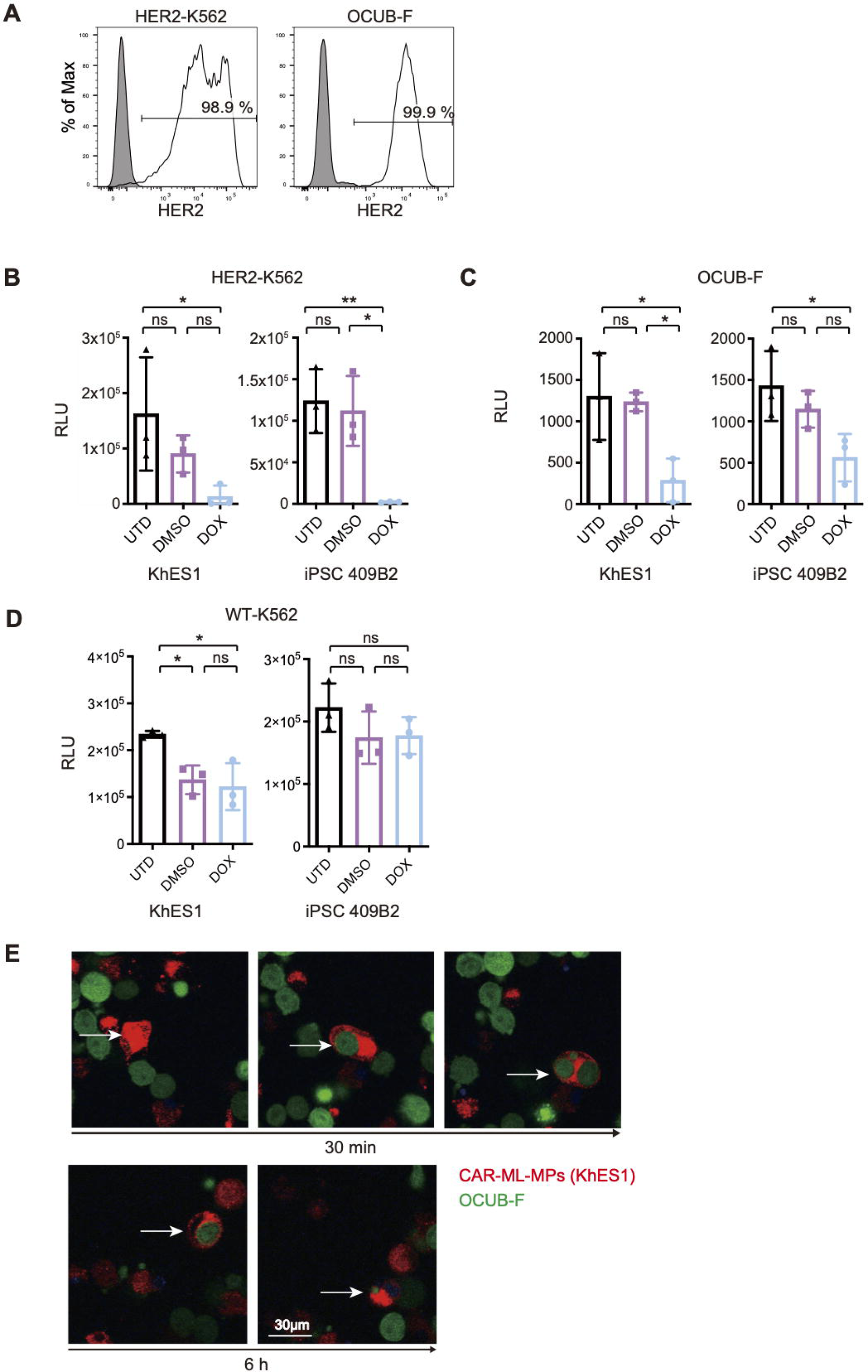
Anti-tumor ability of CAR-ML-MPs *in vitro*. (A) Representative histograms showing HER2 expression in HER2-K562 and OCUB-F. (B, C) Cytotoxicity assay of CAR-ML-MPs derived from KhES1 and iPSC 409B2 strains against HER2-K562 (B) and OCUB-F (C). Effector-to-target ratio were 5:1 to HER2-K562 and 3:1 to OCUB-F, respectively (n = 3, independent experiment). UTD indicates untreated group. DMSO and DOX represent conditions treated without and with induction of CAR expression, respectively. RLU: relative light unit. Dunnett’s tests were performed as an *a priori* multiple comparison. *P < 0.05, **P < 0.01. Error bars represent mean ± standard deviation (SD). (D) Cytotoxicity assay of CAR-ML-MPs against wild type K562. (E) Results of time-lapse imaging of co-culture of CAR-ML-MPs (KhES1 strain) and OCUB-F (E/T ratio 1:1) after 30 minutes and 6 hours. Representative ML-MP cell (white arrow) showed phagocytosis of OCUB-F. See Video S1.

### CAR-ML-MPs have anti-tumor ability *in vivo* but diminish within a week

Subsequently, we validated the *in vivo* cytotoxicity of CAR-ML-MPs. We utilized a tumor-bearing mouse model, in which Luc-HER2-K562 cells, expressing both Luc and HER2, were subcutaneously implanted into immunodeficient NOD/Shi-scid, IL-2Rgnull (NOG) mice (Figure 3A). A single subcutaneous dose of CAR-ML-MPs administered concurrently with the tumor markedly reduced the tumor mass (Figures 3B and 3C) and prolonged the survival of tumor-bearing mice (Figure 3D). The extended survival observed in the DOX-treated group with induced CAR expression unequivocally demonstrates the significant contribution of CAR to the augmentation of anti-tumor effects on the surface of ML-MPs. Furthermore, even in the absence of CAR expression, the survival of mice was statistically significantly prolonged, indicating that some degree of antitumor activity is inherent in ML-derived macrophages (ML-MPs) (Figures 3C and 3D). However, ultimately, mice in all groups failed to prevent tumor enlargement and succumbed to mortality. This suggests a duration limitation in the therapeutic efficacy using ML-MPs, irrespective of CAR expression. Results from a separate experiment involving the administration of luciferase-expressing ML-MPs alone (Figure 3E) supported this notion, as ML-MPs decreased by up to 10% on day 3 following subcutaneous injection and became undetectable by day 7 (Figures 3F and 3G).

**Figure 3.**
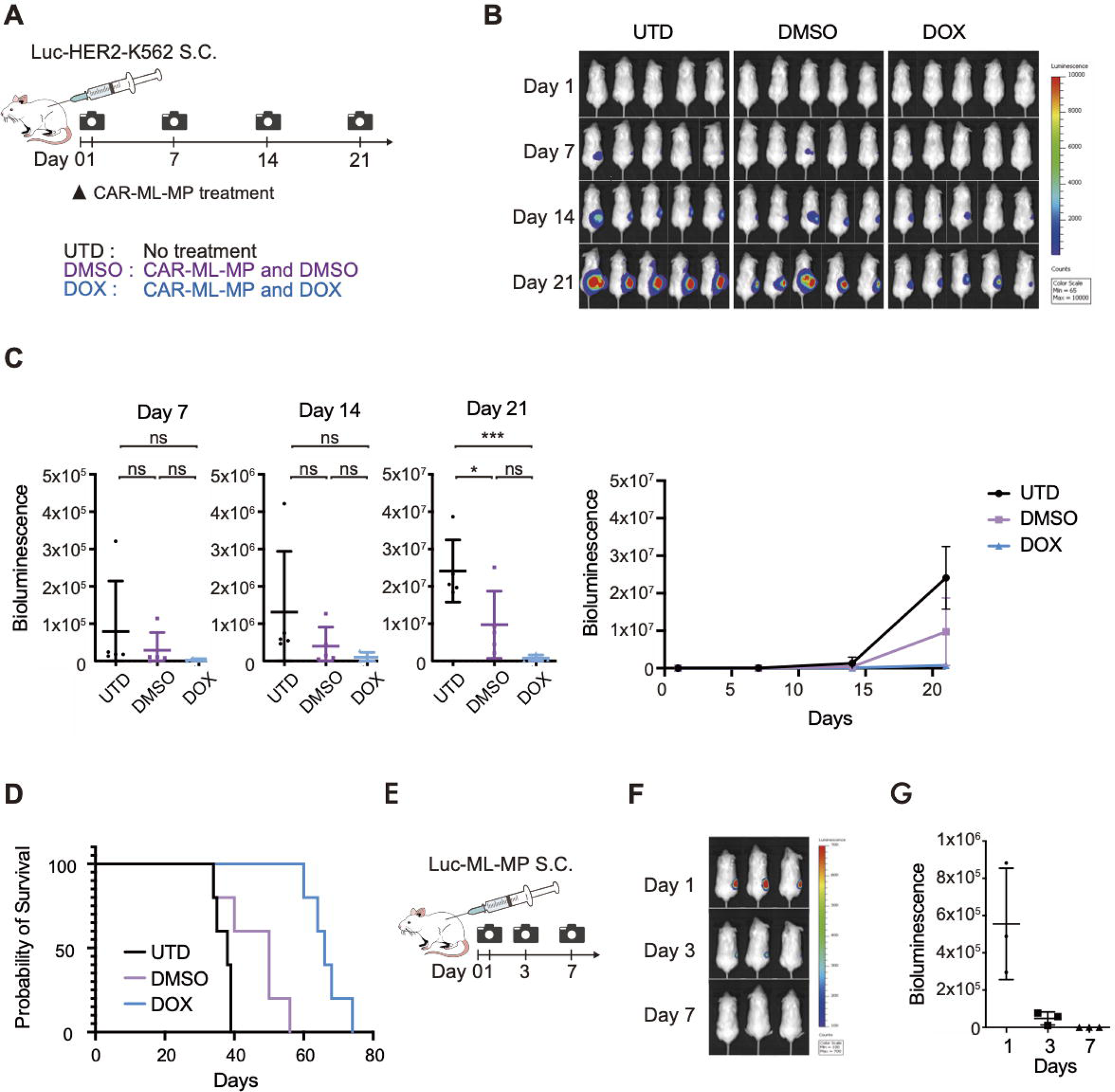
Anti-tumor activity of single administration regimen *in vivo*. (A) Experimental time course of *in vivo* tumor treatment assay. (B, C) Bioluminescence images (B) and quantitative values of bioluminescence (C) from Luc-HER2-K562 cells at the indicated days and conditions. Statistical analysis was performed using *a priori* Dunnett’s multiple comparisons test. UTD indicates untreated group. DMSO and DOX represent conditions treated without and with induction of CAR expression, respectively. *P < 0.05, ***P < 0.001. Error bars represent mean ± standard deviation (SD). (D) Kaplan–Meier plot showing the survival of each group (n = 5 for each group). Statistical analysis was performed with Log-rank (Mantel-Cox) test. (E) Experimental time course of Luc-ML-MP transplantation assay. Bioluminescence images of (F) Luc-ML-MPs and (G) their quantification (n = 3). Error bars represent mean ± standard deviation (SD).

### Repeated administration of CAR-ML-MPs enhances tumor suppression and prolongs survival of tumor-bearing mice

Following the results of the first trial of *in vivo* treatment, we finally examined whether repeated administration of CAR-ML-MPs could enhance the anti-tumor effect compared to a single dose (Figures 4A and 4B). The results demonstrated that an additional six days of repeated intra-tumor administration, with or without CAR expression, significantly suppressed tumor growth and prolonged survival compared to the single treatment group (Figures 4C and 4D). In a separate experiment, one week after continuous CAR-ML-MP administration, no signal was detected, similar to the single administration scenario (Figures 4E-4G). Therefore, continuous administration itself is unlikely to be harmful.

**Figure 4.**
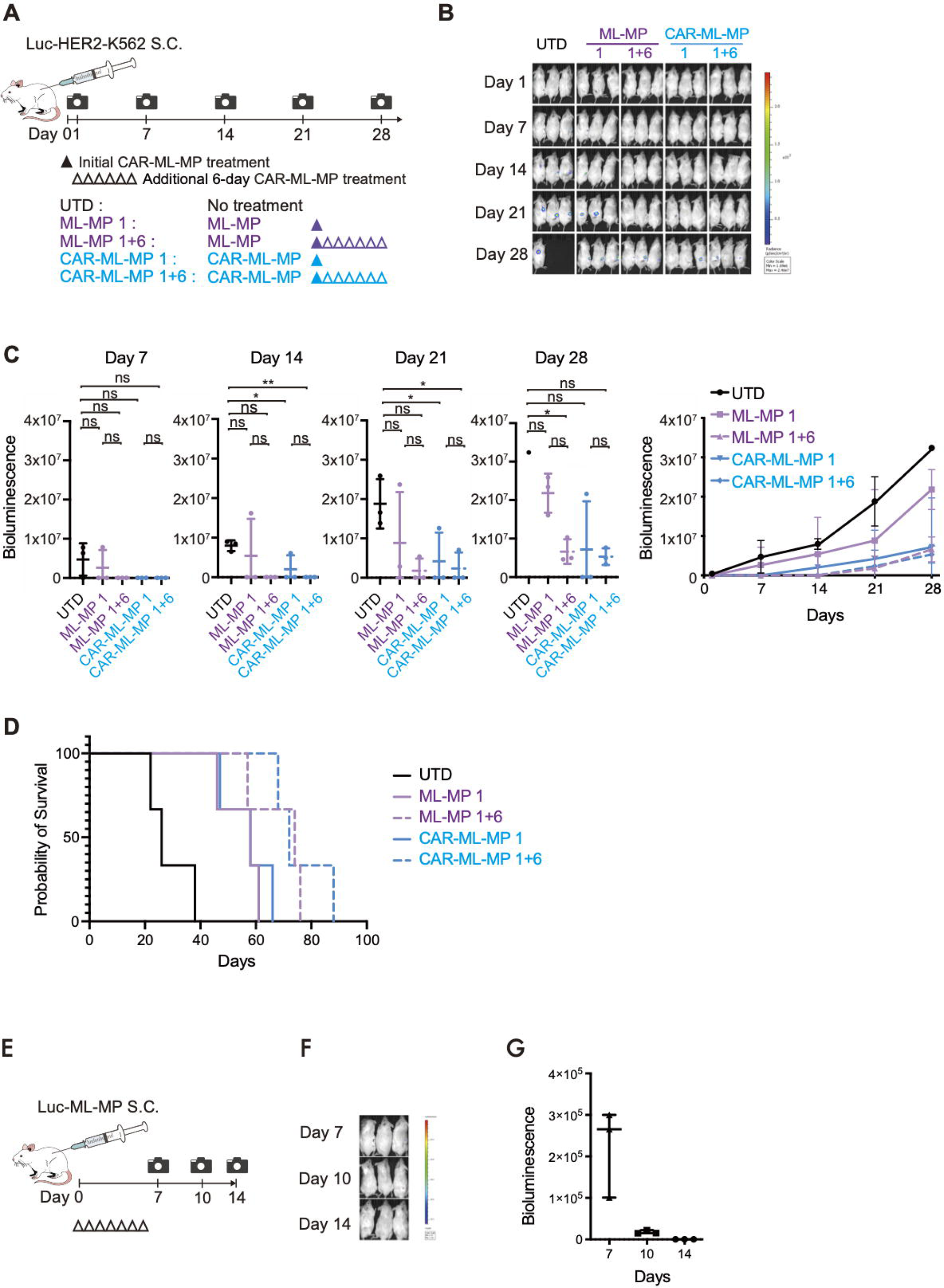
Anti-tumor activity of sequential administration regimen *in vivo*. (A) Experimental time course of *in vivo* assays comparing single and consecutive tumor treatments. (B) Bioluminescence images and (C) quantitative values of bioluminescence from Luc-HER2-K562 cells at the indicated days and conditions. Statistical analysis was performed using *a priori* Dunnett’s multiple comparisons test. UTD indicates untreated group; 1 and 1+6 indicate single treatment alone and with additional treatment, respectively. *P < 0.05, **P < 0.01. Error bars represent mean ± standard deviation (SD). (D) Kaplan–Meier plot showing the survival of each group (n = 3 for each group). Statistical analysis was performed with Log-rank (Mantel–Cox) test. (E) Experimental time course of repeated Luc-ML-MP transplantation assay. Bioluminescence images of (F) Luc-ML-MPs and (G) their quantification (n = 3). Error bars represent mean ± standard deviation (SD).

## Discussion

The current study demonstrates for the first time that ML-MPs possess the advantages of labor- and time-saving preparation and exhibit anti-tumor activity both in vitro and in vivo. By proliferating cells sufficiently in the ML state before differentiating them into macrophages, it is possible to prepare large quantities of macrophages easily and continuously. Although there were some differences in the expression profiles among the strains, all examined ML strains could be cultured into ML-MPs. The cytotoxic effects of ML-MPs were enhanced by CAR expression and repeated administration, suggesting that CAR-ML-MPs are suitable off-the-shelf sources for more cost-effective and efficient adoptive cell therapy (ACT). Due to experimental and interindividual variations among mice, especially in the absence of CAR, it would be necessary to conduct larger-scale studies to evaluate the efficacy of single-dose treatments accurately. However, the results of repeated administration demonstrated a favorable trend compared to the untreated group, and the combination with CAR further strengthened this effect, delaying tumor growth.

Macrophages phagocytose tumor cells and present tumor antigens^6^, which could confer long-term and potent tumor immunity. Although an exact comparison of efficacy is difficult, we confirmed that CAR-ML-MPs phagocytosed live tumor cells in vitro via time-lapse imaging. Moreover, repeated administration prolonged the duration of the effect; in Figure 4, tumor re-growth in vivo was observed from day 7 after a single dose of ML-MPs, whereas it was observed 7 days later (day 14) for CAR-ML-MPs. This is consistent with the results shown in Figure 2, indicating the stronger cytotoxic activity of CAR-ML-MPs. It is possible that the single administration of CAR-ML-MPs alone was sufficient to reduce the residual tumor volume enough to slow down tumor re-growth, resulting in no significant difference between single and repeated administration.

As CAR-ML-MPs in this study express HLA-DR (MHC class II antigen)^15^ and ML-derived dendritic cells were previously reported to activate HLA-compatible T cells, CAR-ML-MPs may contribute to the activation of T-cell cancer immunity in addition to direct phagocytosis. According to previous reports, to obtain 1 x 10^9^ monocytes from iPS cells, 500-5,000 undifferentiated iPS cell colonies need to be cultured for 3-4 weeks to induce differentiation. In contrast, the MLs used in this study, once established, can be inexpensively cultured for maintenance and rapid proliferation, and can also be cryopreserved. This means that the cost of medium, cytokines, and other reagents, as well as the human and equipment resources required for cell preparation, can be significantly reduced.

MHC compatibility should be considered when performing allogeneic ML-MP transplantation. In Japan, iPS cells have been established from MHC-homo donors for allogeneic transplantation^16^. MHC-compatible transplantation should not only reduce immune rejection, but also activate antigen-specific T cells via antigen presentation from allogeneic cells. Although evaluation of the antigen presentation ability of CAR-ML-MPs and adoptive immunity induction should be verified in more detail *in vitro* and *in vivo*, the use of iPS cells produced for allogeneic transplantation could lead to more affordable and stable adoptive cellular therapy.

A current limitation of this study is the potential safety concerns surrounding MLs, as the transduced proto-oncogenes that may induce tumorigenesis. MLs rapidly diminish in the mouse body and become almost undetectable 48 h after transplantation^17^. Consistent with this study, our results showed that ML-MPs decreased rapidly in mice and became undetectable within seven days, even with repeated administration. Thus, in the current study, the likelihood of tumorigenesis in MLs and ML-MPs was considered low. Nevertheless, administering adoptive cellular immunotherapy to patients remains difficult without further safety assurance. Because this study focuses on a nonclinical proof-of-concept, further safety validation was not performed at this time. Future clinical trials, including the establishment of MLs without proto-oncogenes, irradiation to stop the cell cycle, and induction of suicide genes such as HSV-TK/GCV or inducible caspase-9 system^18^ are needed to translate the results of this study to the bedside.

A second limitation of this study is that although the potential of CAR-ML-MP was demonstrated, its cytotoxic efficacy was only assessed in HER2-expressing tumor cells. Despite the widely recognized efficacy of CAR in combination with cell therapy for B-cell lineage malignancies, the efficacy against other tumors, including solid tumors, has not yet been fully recognized. Therefore, more tumor cell models will need to be investigated in the future to demonstrate the applicability of this approach to a wide range of tumor therapies. Such trials will require not only the identification of optimal antigens that are central targets for anti-tumor activity, but also the modification of the immune cells themselves to increase their ability to infiltrate tumors and their duration of efficacy, as well as the improvement of treatment regimens. The combination of CAR and ML-MPs, as demonstrated in this study, needs to be explored in more depth in this context. When such cells are established, repeated-dose therapy, with the technical advantage of easier preparation, may provide an answer to the conflicting demands for safety and prolonged anti-tumor efficacy. Regimen optimization, which may involve combining other cell transplantation therapies or immune checkpoint therapies, such as anti-CD47 and anti-PD-1 antibody, holds the potential to improve anti-tumor efficacy and present a novel treatment option for cancer.

## Acknowledgements

We thank Dr. Yasushi Uemura (National Cancer Center, Tokyo, Japan) for kindly providing Luc-K562, Dr. Cliona M. Rooney (Baylor College of Medicine) for kindly providing a plasmid vector encoding anti-HER2 CAR construct, Dr. Akihiro Ikenaka for supporting time-lapse imaging, and Ms. Harumi Watanabe for providing administrative assistance. This work was supported by the grant for the Core Center for iPS Cell Research of the Research Center Network for Realization of Regenerative Medicine (JP21bm0104001) from the Japan Agency for Medical Research and Development (AMED) to M.K.S., the Program for Intractable Diseases Research utilizing Disease-specific iPSC cells (JP21bm0804004) from AMED to M.K.S., the Core Center for Regenerative Medicine and Cell and Gene Therapy (JP23bm1323001) from AMED to M.K.S., a grant from the Terumo Life Science Foundation to M.K.S., and a grant from the iPS Cell Research Fund to M.K.S.

## Author contribution

Y.A. and A.N. designed and performed experiments and analyzed data; T.K. performed experiments and analyzed data; S.Y. and Y.N. provided CAR construct; M.K.S. supervised the project and experimental design; and A.Y., A.N. and M.K.S. wrote the paper.

## Supplemental Movie

**Supplemental Movie 1. Phagocytosis of tumor cells by ML-MPs.**

